# Contribution of murine strain background to Na^+^ reabsorption in the kidney

**DOI:** 10.64898/2026.03.17.712246

**Authors:** Sarah Christine M. Whelan, Stephanie M. Mutchler, Sophia Mitton-Fry, Leeann Parsi, Sri Balaji, Thomas R. Kleyman, Shujie Shi

## Abstract

Kidneys play an essential role in balancing fluid and electrolyte levels. Two mouse strains, C57Bl/6 and 129S2/SV, are routinely used to study renal physiology in laboratory settings, and prior observations suggest that significant differences in salt and water handling exist between them. This study aims to further establish the sources of these observed differences at both expressional and functional levels, in male and female mice. At baseline, male 129S2/SV mice displayed decreased Na^+^ and increased K^+^ plasma concentrations compared to C57Bl/6 males, while no statistical differences were observed between female mice. Interestingly, 129S2/SV male mice had lower glomerular density than C57Bl/6 males. Immunoblotting shows that 129S2/SV mice of both sexes had increased expression of NHE3 and NKCC2 compared to their C57Bl/6 counterparts. Both total and phosphorylated NCC were more abundant in female mice as compared to males, indicating sexual dimorphism. Furthermore, 129S2/SV females had higher expression of total and phosphorylated NCC compared to C57Bl/6 females. In contrast, the expression of SGLT2, ENaC subunits, and Na^+^/K^+^-ATPase were comparable between C57Bl/6 and 129S2/SV mice of both sexes. When challenged with diuretics intended to block NKCC2, NCC or ENaC, 129S2/SV male mice responded with a smaller diuresis and natriuresis than their C57Bl/6 counterparts. Taken together, our data suggest that differential expression of key Na^+^ transporters along the nephron contributes to differences in Na^+^/K^+^ homeostasis between these two mouse strains.

**NEW & NOTEWORTHY:** We assessed the influence of genetic background on the expression of key Na^+^ transporters along the nephron in two commonly used inbred mouse strains, C57Bl/6 and 129S2/SV. We found that the kidney expression of NHE3, NKCC2, and NCC are strain dependent. Additionally, murine strain significantly contributes to the diuretic responses induced by hydrochlorothiazide, amiloride, and furosemide.

## INTRODUCTION

The majority of glomeruli-filtered serum Na^+^ is reabsorbed at three sites along the nephron (1): the proximal tubule (PT), the thick ascending limb (TAL) of the loop of Henle, and the distal nephron which includes the distal convoluted tubule (DCT), the connecting tubule (CNT), and the collecting duct (CD). Approximately two-thirds of tubular Na^+^ reabsorption occurs in PTs and is coupled with the secretion of H^+^ through a Na^+^/H^+^ exchanger (NHE3), reabsorption of glucose via Na^+^-glucose co-transporters (SGLT1 and SGLT2), as well as other Na^+^-coupled co-transporters (2-7). The Na^+^-K^+^-Cl^-^ co-transporter (NKCC2), expressed in the TAL, is responsible for another 15-25% of tubular Na^+^ reabsorption and is inhibited by loop diuretics, such as furosemide (8). The Na^+^-Cl^-^ co-transporter (NCC), a target of thiazide diuretics, reabsorbs ∼5-10% of filtered Na^+^ in the DCT (9). The final site of Na^+^ reabsorption occurs in the late DCT and CNT/CD via the epithelial Na^+^ channel (ENaC), providing the driving force for K^+^ secretion (10-13). The K^+^-sparing diuretic amiloride and its derivatives inhibit ENaC by directly binding within the channel’s pore region (14, 15). Patients with loss-of-function mutations in genes encoding several of these transporters (e.g. *Slc9a3, Slc12a1, Slc12a3, Scnn1a, Scnn1b* or *Scnn1g*) exhibit defective Na^+^ reabsorption, leading to aberrant extracellular volume and blood pressure control (16-18).

C57Bl/6 is one of the most well-established experimental inbred mouse strains and serves as the background for several genetic models used to study renal (patho)physiology. In recent years, the 129S2/SV strain has gained popularity as an experimental model of salt-sensitive hypertension due to its extra copy of the *Renin* gene and enhanced sensitivity to high salt diets, as compared to C57Bl/6 mice (19). Furthermore, 129S2/SV mice have lower GFR and are more sensitive to renal ischemia-reperfusion injuries (20, 21). Despite clear disparities in renal phenotypes between these two strains, their differences in salt and water handling have not been carefully assessed. The present study compares the abundance and post-translation modifications (phosphorylation, proteolysis) of the major renal Na^+^ transporters and channels between C57Bl/6 and 129S2/SV mice. Additionally, NKCC2, NCC, and ENaC function was compared between the two strains by challenging the mice with diuretics.

## MATERIALS AND METHODS

### Animals and Tissues

All animal protocols used in the present study adhere to the NIH Guide for the Care and Use of Laboratory Animals and were approved by the University of Pittsburgh IACUC (protocol 23124129). Adult male and female mice (>10 weeks old) of C57Bl/6NCr1 (Charles River #027) or 129S2/SVPasCr1 (Charles River #476) lineage were housed in a temperature-controlled, 12:12 h light/dark photocycle facility with free access to water and standard pellet chow. Blood was collected via cardiac puncture for iSTAT Chem8^+^ analysis (Abbott Laboratories #09P31-26) following isoflurane anesthesia. Animals were sacrificed by exsanguination and confirmed by cervical dislocation. Kidneys were collected for further analysis.

### Histology

Whole kidneys were coronally cut and fixed in 10% formalin. Tissues were paraffin embedded and sectioned at 3μm. Hematoxylin and Eosin (H&E) staining was completed by University of Pittsburgh’s Research Histology Biospecimen Core. Sections were imaged on a Leica widefield microscope (DM6000) and stitched together using ImageJ (NIH, RRID: SCR_003070). Glomeruli counting within the cortex was performed blinded and glomerular density was presented as number of glomeruli per mm^2^ cortex.

### Western Immunoblot

Kidneys were quartered and lysed in ice-cold CellLytic buffer (Sigma #C3228) supplemented with 1% protease and phosphatase inhibitors (Sigma #539134, #524625). Lysates were tumbled for 30 minutes at 4°C and centrifuged at 4,000g for 10 minutes. The supernatant was spun at 100,000g for an additional 60 minutes to enrich membrane-bound proteins. The pellet was resuspended in 300 μL lysis buffer and concentration was determined using a BCA assay (ThermoFisher #23225). 60 μg of protein was separated in Criterion 4-15% TGX stain-free gels (BioRad #5678085) under reducing conditions (5% β-mercaptoethanol; BioRad #1610710), transferred to nitrocellulose membranes (Millipore #HAWP04700), blocked with 5% milk for 1h at room temperature, and incubated with primary antibody (see Table 1) at 4° overnight with gentle rocking. Following 2h incubation with HRP-conjugated secondary antibody (see Table 1) at room temperature, membranes were developed with BioRad Clarity Substrates (#1705061, #1705062) and imaged on a ChemiDoc. Bands were quantified using ImageLab densitometry software and normalized to total protein loading obtained with stain-free technology.

**Table 1.** Primary and secondary antibodies used in western blot analysis.

| **Antibody** | **Source** | **Catalog #** | **PRID** | **Dilution** |
| --- | --- | --- | --- | --- |
| Rabbit anti-SGLT2 | ProteinTech | 24654-1-AP | AB_2750601 | 1:2000 |
| Mouse anti-NHE3 | Stressmarq | SPC400 | AB_10643557 | 1:2000 |
| Rabbit anti-NKCC2 | Stressmarq | SPC401 | AB_10640877 | 1:2000 |
| Rabbit anti-NCC | Millipore | AB3553 | AB_571116 | 1:2000 |
| Rabbit anti-pNCC(T<sup>53</sup>) | Phosphosolutions | P1311-53 | AB_2650477 | 1:2000 |
| Rabbit anti-αENaC | Dr. J. Loffing | N/A | N/A | 1:2000 |
| Rabbit anti-βENaC | Dr. L. Palmer | N/A | N/A | 1:2000 |
| Rabbit anti-γENaC | Stressmarq | SPC-405 | AB_10640369 | 1:3000 |
| Mouse anti-NKAα1 | Developmental Studies Hybridoma Bank | a5 | AB_21665869 | 1:1000 |
| Donkey anti-rabbit HRP | Jackson ImmunoResearch | 711-035-152 | AB_10015282 | 1:5000 |
| m-IgG Fc BP-HRP | Santa Cruz | sc-525409 | AB_3101828 | 1:5000 |

### Metabolic Cages

Mice were individually housed in wire-bottom metabolic cages for urine collection with two days acclimation and free access to standard-extruded chow (0.4% NaCl). Intraperitoneal sham injections (100 μl sterile saline) were administered at 9 PM, with urines collected 3h post-injection. Following a 24-hour washout, mice were injected with 100 μl saline containing 5 mg/kg amiloride (Merck #MK685), 25 mg/kg hydrochlorothiazide (HCTZ, Sigma #H2910), or 30 mg/kg furosemide (Sigma #F4381) with urine collected 3h post-injection. Body weight and urine volume were documented daily. Urinary [Na^+^], [K^+^], and [Cl^-^] were measured using the EasyLyte analyzer (Medica Corporation). Urinary excretion of Na^+^ (U_Na_V), K^+^ (U_K_V), and Cl^−^ (U_Cl_V) were normalized to body weight (BW) and shown as µmol/gBW. Diuretic-induced changes (Δ) in urine volume, U_Na_V, U_K_V, or U_Cl_V were calculated by subtracting the sham injection values from the same animal.

### Statistics

Data were analyzed using Prism software and presented as mean ± SD. Main effects were analyzed with ordinary two-way ANOVA. Regardless of the significance of main effects, pre-specified effects (male C57Bl/6 x male 129S2/SV, male C57Bl/6 x female C57Bl/6, male 129S2/SV x female 129S2/SV, female C57Bl/6 x female 129S2/SV) were analyzed using Sidak’s multiple comparisons test because they directly address our biological questions regarding strain differences within each sex.

## RESULTS

We first examined blood chemistries in mice of the two background strains. When compared to C57Bl/6 male mice, 129S2/SV males had a significantly lower blood [Na^+^] and higher blood [K^+^], as well as a higher [Cl^-^] that approached significance. No significant differences in blood chemistry were detected between female mice of the two strains (Figure 1, A-C). We also measured kidney weight and body weight in age-matched male and female C57Bl/6 and 129S2/SV mice. As expected, male mice had higher body weights and wet kidney weights than female mice of the same background strain (Figure 1, D-E). We performed H&E staining to compare kidney histology and found no gross differences between the two strains (Figure 1G). However, glomeruli counting in the renal cortex demonstrated that 129S2/SV male mice have lower glomerular density compared to C57Bl/6 males (Figure 1H). Again, females showed no significant difference.

**Figure 1.**
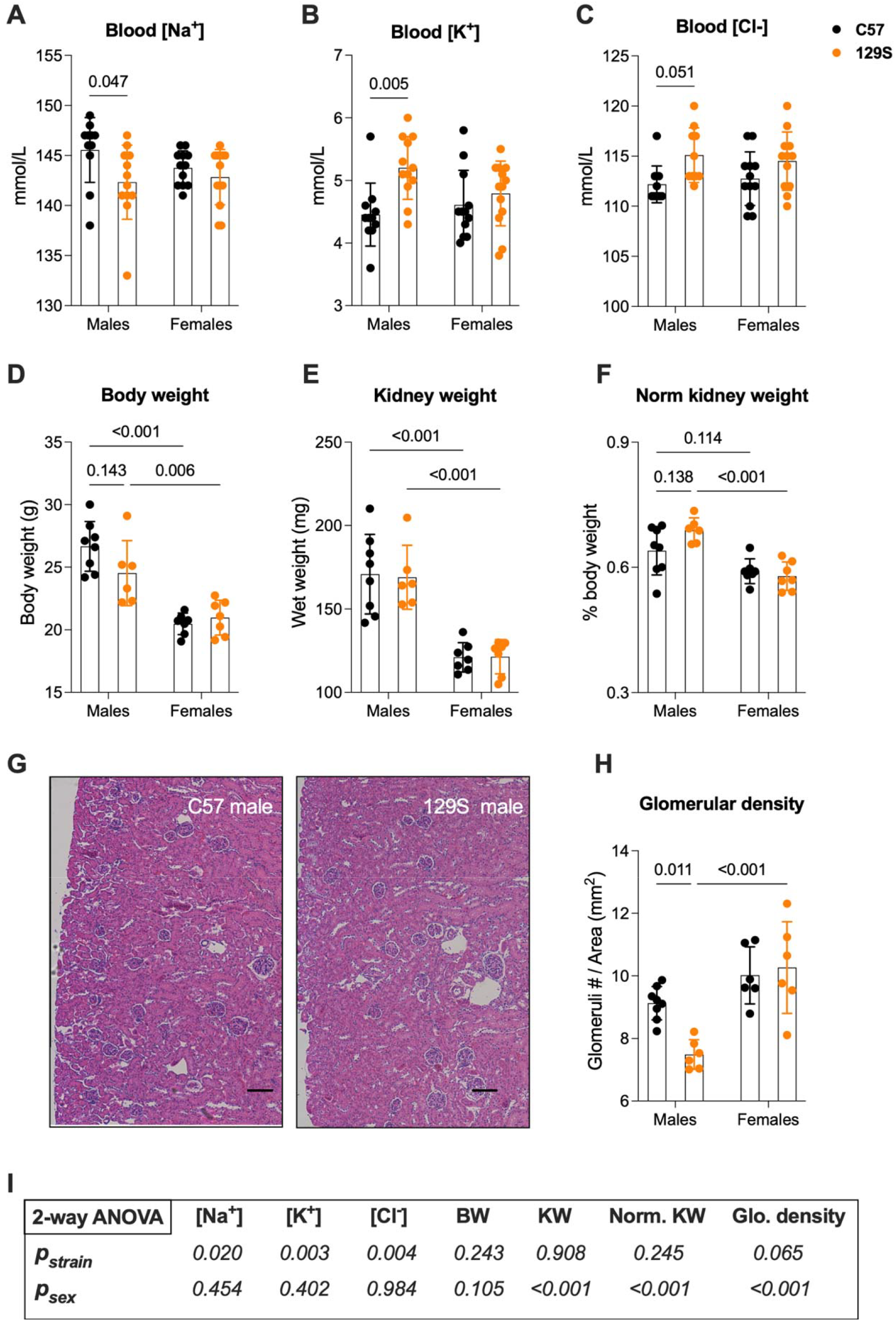
Baseline blood electrolytes and kidney morphology in age-matched C57 and 129S mice. Adult male and female mice were maintained on regular chow with free access to water. **(A-C)** iSTAT Chem8^+^ analysis of blood electrolytes are presented. Body weight **(D)** and wet kidney weight **(E)** of individual mice were documented to calculate normalized kidney weight **(F). (H)** Glomerular density was determined by counting the number of glomeruli per mm^2^ renal cortical area using H&E images as shown in **(G)**, scale bars = 50 μm. Statistical comparisons between strains and sexes were performed by two-way ANOVA, followed by Sidak’s multiple comparison test to control for family-wise error rate. *P* values of < 0.2 are shown. The main effect results of the 2-way ANOVA are reported in the table **(I)**.

Next, we probed for the abundance of major Na^+^ transporters and channels at sites of reabsorption along the nephron, including NHE3 and SGLT2 within the PT; NKCC2 in the TAL; NCC in the DCT; and α, β, γ-subunits of ENaC in the late DCT, CNT and CD (Figure 2). The α subunit of Na^+^/K^+^ ATPase (NKAα1) that is ubiquitously expressed along the nephron was also measured (Figure 2). 129S2/SV female mice had lower abundance of SGLT2 compared to both 129S2/SV males and C57Bl/6 females. 129S2/SV mice of both sexes displayed higher abundance of NHE3 and NKCC2 compared to C57Bl/6 mice. Expression of total NCC and phosphorylated NCC at T^53^, a phosphorylation site associated with NCC activation (22), was greater in female mice of both strains, with 129S2/SV females displaying even greater expression than their C57Bl/6 counterparts. Differences in ENaC subunit expression were not as straightforward. While there were no significant differences in full-length αENaC, female 129S2/SV mice had increased abundance of cleaved αENaC as compared to C57Bl/6 females. In contrast, male C57Bl/6 mice had increased abundance of cleaved αENaC as compared to 129S2/SV males. βENaC expression was lower in 129S2/SV females versus 129S2/SV males, with no difference detected between strains of the same sex. The expression of full-length and cleaved γENaC was comparable between sexes and strains. NKAα1 expression was also similar between sexes and strains. Overall, background strain affects renal expression of NHE3, NKCC2, NCC and pNCC, while sexual dimorphism was displayed in NHE3, NCC, pNCC, cleaved αENaC, βENaC and NKAα1 abundance (Figure 2C).

**Figure 2.**
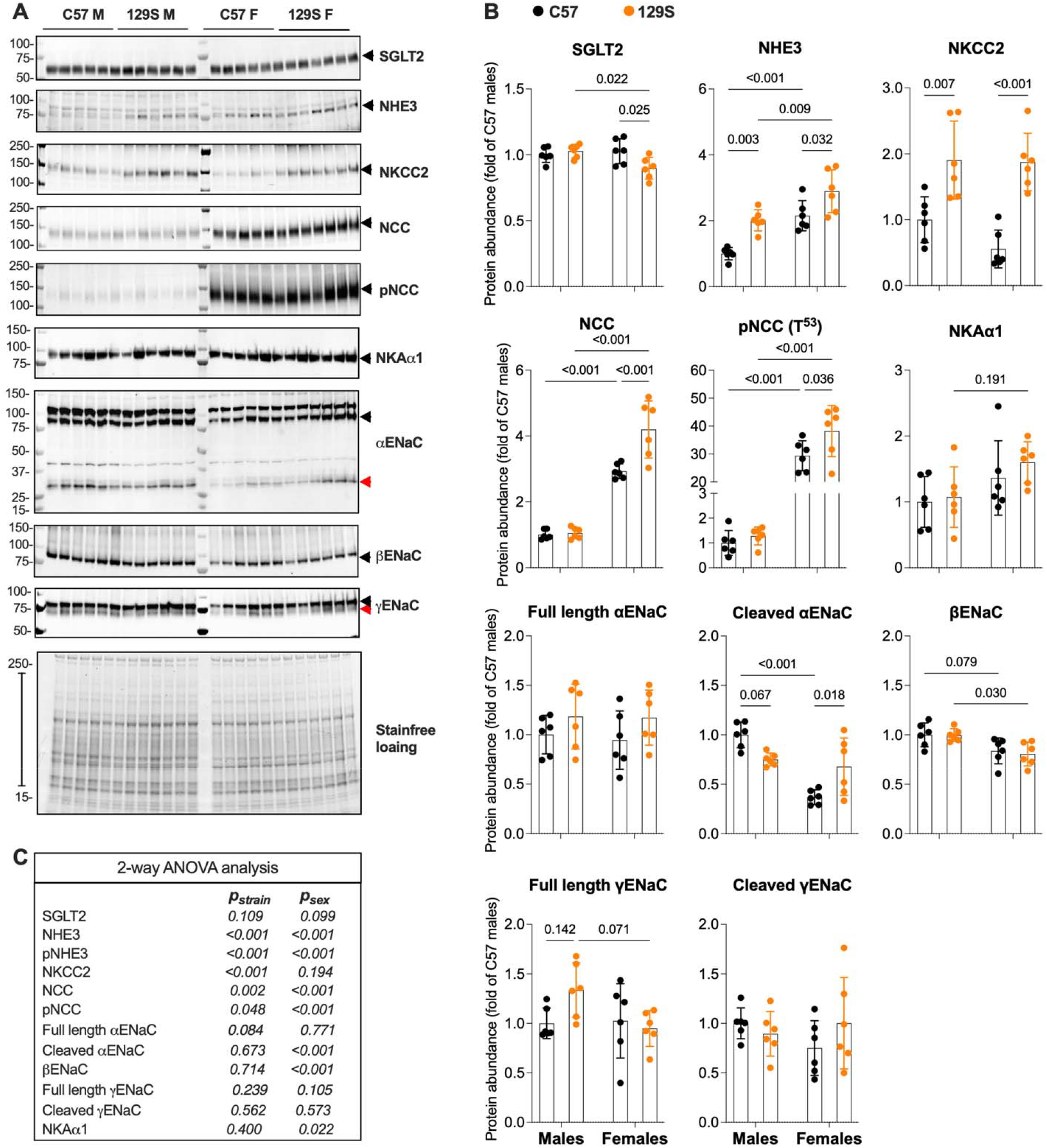
Immunoblotting analysis of Na^+^ transport along the nephron. Kidneys were harvested from C57 (•) and 129S 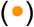 adult male and female mice, homogenized, and centrifuged to collect membrane enriched fractions. (**A**) Representative blots showing SGLT2, NHE3, NKCC2, NCC, pNCC, ENaC α, β or γ subunit and NKAα1. Arrow heads indicate bands utilized for quantification, with red arrow heads indicating the cleaved α and γ ENaC subunits. Markers of molecular weight (kDa) are shown on the left. (**B**) The abundance of each protein target was normalized to the total protein in each corresponding sample (estimated with strain-free images) and is shown as fold of C57 male mice. Statistical comparisons were analyzed using two-way ANOVA, followed by a Sidak’s multiple comparison test to control for family-wise error rate, with *P* values < 0.2 shown for each comparison. The main effects results of the 2-way ANOVA are shown in the table **(C)**.

Lastly, we examined the function of NKCC2, NCC and ENaC by measuring the diuretic and natriuretic response to a single-dose IP injection of furosemide, HCTZ, or amiloride, respectively. All three inhibitors increased diuresis and natriuresis compared to that of the saline injection, confirming the targeted Na^+^ reabsorption route was sufficiently blocked (Figure 3). Urine volume changes in response to NKCC2 inhibition with furosemide were similar between the two mouse strains and between both sexes. Furosemide-induced natriuresis was significantly higher in male C57Bl/6 mice than female C57Bl/6 mice, but comparable between the two strains (Figure 3A). NCC inhibition with HCTZ evoked significantly higher urine output and natriuresis in C57Bl/6 males than 129S2/SV males, with no difference detected between female mice of the two strains. Kaliuresis in response to HCTZ was also influenced by background strain, although not reaching statistical significance among the four groups of mice (Figure 3B). Both male and female C57Bl/6 mice exhibited a greater natriuretic response to amiloride, compared to their 129S2/SV counterparts. Kaliuresis in response to amiloride was similar among all four groups of mice (Figure 3C). No significant difference was observed in furosemide-, HCTZ-, or amiloride-induced chloruresis between the two strains or sexes (Figure 3). Overall, background strain was a major contributor to NKCC2 and NCC activity only in male mice, while ENaC activity was affected by background strain in both male and female mice (Figure 3D).

**Figure 3.**
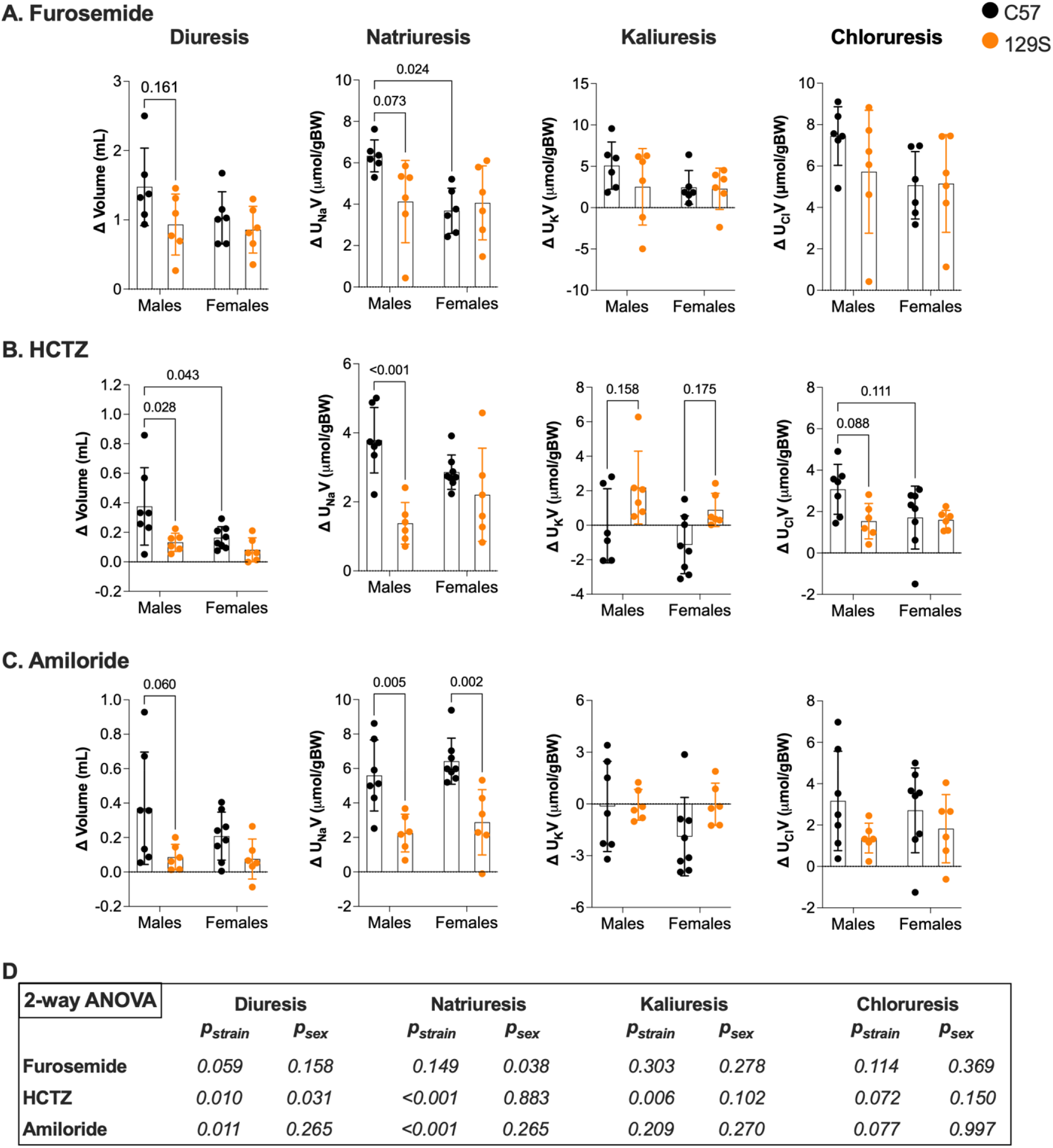
Urine output following diuretic challenges. Mice were given a single IP injection of diuretic (5mg/kg amiloride, 25mg/kg hydrochlorothiazide, or 30mg/kg furosemide) 24h following administration of sham injection. Urine samples were collected 3h post-injections. Diuretic induced changes in urine volume and urinary excretion of Na^+^ (ΔU_Na_V), K^+^ (ΔU_K_V), or Cl^−^ (ΔU_Cl_V) from individual mice are shown for furosemide **(A)**, HCTZ **(B)**, and amiloride **(C)**. Statistical comparisons were analyzed using two-way ANOVA, followed by a Sidak’s multiple comparison test to control for family-wise error rate, with *P* values < 0.2 shown for each comparison. The main effects results of the 2-way ANOVA are shown in the table **(D)**.

## DISCUSSION

Sexual dimorphism in renal electrolyte transport is well known (23-27), but the influence of mouse genetic background on salt and water handling has not been updated in recent years (28-31). In the present study, we confirmed the importance of sex and assessed the contribution of background strain by examining the expression and function of major renal Na^+^ reabsorption pathways in males and females of two commonly used inbred mouse strains, C57Bl/6 and 129S2/SV. When compared to their C57Bl/6 counterparts, male and female 129S2/SV mice had higher expression of NHE3 and NKCC2. Additionally, female 129S2/SV mice had higher total and phosphorylated NCC. Surprisingly, 129S2/SV mice exhibited smaller diuretic and natriuretic responses to both furosemide and HCTZ than C57Bl/6 mice. Despite the sex-specific variations in ENaC subunit cleavage (reduced α subunit cleavage in male 129S2/SV compared to male C57Bl/6 but enhanced α subunit cleavage in female 129S2/SV compared to female C57Bl/6), amiloride-induced diuresis and/or natriuresis are lower in both male and female 129S2/SV mice compared to their C57Bl/6 counterparts. The reduced natriuretic response to amiloride in 129S2/SV mice, compared to C57Bl/6 mice, is consistent with the differences in ENaC open probability (P_o_) that exist between the two strains. We have previously observed that the 8 pS channel (ENaC with Li^+^ as the charge carrier) has lower average P_o_ (equal to ∼0.2 or less) in 129S2/SV mice, compared to the reported ENaC P_o_ in C57Bl/6 mice (normally ∼0.4 or greater) when mice are on a control or a low salt diet (32-39).

Overall, our immunoblotting analyses suggest that the Na^+^ reabsorption machinery in the PT (NHE3), loop of Henle (NKCC2) and DCT (NCC) are upregulated in 129S2/SV mice (Figure 2). Surprisingly, blocking NKCC2 or NCC with furosemide or HCTZ was associated with reduced natriuresis in 129S2/SV mice compared with C57Bl/6 mice (Figure 3). It is reasonable to speculate that the greater expression of NHE3 in 129S2/SV mice allows a larger portion of Na^+^ reabsorption to occur in the PT, minimizing the distal delivery of Na^+^. This would explain the discrepancies between transporter abundances and diuretic responses in the two examined murine strains. However, our observation of differences in glomerular density and blood electrolytes between male mice of the two strains adds further complexity to the story (Figure 1). While this study is not a complete survey of channels and transporters that participate in Na^+^ reabsorption along the nephron, it provides an update on the effect of mouse background strain on renal Na^+^ reabsorption and adds to a growing body of literature that demonstrates genetic background can affect (patho)physiology and even behaviors. Taken into consideration alongside studies showing the effect of strain differences in susceptibility to renal injury and fibrosis (20, 40-43), anatomical features in the kidney, and GFR variability (44), our data serve as a reminder that strain matters and proper reporting of sub-strain and vendor is critical to scientific rigor and reproducibility.

## DATA AVAILABILITY

All original immunoblot images presented in the manuscript have been archived and are fully available to the journal upon reasonable request.

## AKNOWLEGEMENT

We thank the Dr. J. Loffing for providing the anti-αENaC antibody and Dr. L. Palmer for sharing the anti-βENaC antibody.

## AUTHOR CONTRIBUTIONS

S.M.M. and S.S. conceived and designed research; S.C.M.W., S.M.M., S.M., L.P., K.G.S. and S.B. performed experiments; S.C.M.W., S.M.M., S.M., L.P. analyzed data; S.C.M.W., S.M.M., and S.S. interpreted results; S.C.M.W. and S.S. prepared figures; S.C.M.W. S.M.M and S.S. drafted manuscript; S.C.M.W., S.M.M., T.R.K., and S.S. edited and revised manuscript; S.C.M.W., S.M.M., S.M., L.P., K.G.S., S.B., T.R.K. and S.S. approved final version of manuscript.

